# Acibenzolar-S-methyl activates stomatal-based defense systemically in Japanese radish by inducing peroxidase-dependent reactive oxygen species production

**DOI:** 10.1101/2020.05.24.113878

**Authors:** Nanami Sakata, Takako Ishiga, Shizuku Taniguchi, Yasuhiro Ishiga

## Abstract

Acibenzolar-S-methyl (ASM) is a well-known plant activator, which is a synthetic analog of salicylic acid (SA). Recently, copper fungicides and antibiotics are major strategies for controlling bacterial diseases. However, resistant strains have already been found. Therefore, there is an increasing demand for sustainable new disease control strategies. We investigated the ASM disease control effect against *Pseudomonas cannabina* pv. *alisalensis* (*Pcal*), which causes bacterial blight on Japanese radish. In this study, we demonstrated that ASM effectively suppressed *Pcal* disease symptom development associated with reduced bacterial populations on Japanese radish leaves. Interestingly, we also demonstrated that ASM activated systemic acquired resistance (SAR), including stomatal-based defense, not only on ASM treated leaves, but also on untreated upper and lower leaves. Reactive oxidative species (ROS) are essential second messengers in stomatal-based defense. We found that ASM induced stomatal closure by inducing ROS production through peroxidase. These results indicate that stomatal closure induced by ASM treatment is effective for preventing *Pcal* pathogen invasion into plants, and in turn reduction of disease development.

## Introduction

Recently, several bacterial pathogens were identified as causal agents of severe worldwide epidemics. *Pseudomonas cannabina* pv. *alisalensis* (*Pcal*) causes bacterial blight of crucifer plants (Brassicaceae plants) (Takikawa and Takahashi, 2014) and inflicts great damage on crucifer production including cabbage, Chinese cabbage, and Japanese radish. On Japanese radish, disease symptoms caused by a newly isolated *Pcal* were observed on leaves and roots (Takikawa and Takahashi, 2014). Discoloration of roots and necrotic leaf lesions have serious implications for the commercial value of Japanese radish. Chemical treatments such as copper fungicides and antibiotics are popular strategies for bacterial disease control. However, *Pcal* strains resistant against these chemicals were found (Takahashi et al., 2013). Therefore, the demand for developing sustainable alternative strategies has been increasing.

Acibenzolar-S-methyl (ASM), a synthetic analog of salicylic acid (SA), has been used to protect crops from several diseases by activating plant defense (Kunz et al., 1997; Oostendorp et al., 2001). We demonstrated that ASM soil drench application induced systemic acquired resistance (SAR) against *Pcal* (Ishiga et al., 2020). SAR processes can be divided into three steps: local immune activation, information relay from local to systemic tissues by mobile signals, and defense activation and priming in systemic tissues (Jung et al., 2009; Shah and Zeier, 2013; Wang et al., 2018). Thus, our previous results implied that mobile signals generated by ASM triggered SAR in untreated leaves. Pre-treating cucumber plant first leaves with ASM protected whole plants from fungal infection with *Colletotrichum orbiculare* (Cools and Ishii, 2002). In cucumber plants, *SAR* genes were rapidly and highly expressed in upper leaves after first leaf treatment with ASM (Cools and Ishii, 2002; Narusaka et al., 1999; Narusaka et al., 2001). However, the ASM-induced SAR mechanism is still unclear. We also demonstrated that ASM protects cabbage plants from *Pcal* by activating stomatal-based defense (Ishiga et al., 2020). Furthermore, ASM induced stomatal-based defense resulted in reduced *Pcal* bacterial populations as well as disease symptoms on cabbage within 4 h after soil drench, and this positive effect is prolonged for at least three weeks (Ishiga et al., unpublished). Additionally, many foliar bacterial pathogens target stomata as an entry site. Therefore, ASM is expected to have a powerful disease control effect. However, whether ASM is effective at suppressing disease development against *Pcal* on other crops (in addition to cabbage) has not been evaluated.

ROS production at the cell surface, which is called oxidative burst, is one of the earliest defenses detectable during pathogen-associated molecular pattern (PAMP)-triggered immunity (PTI) and effector-triggered immunity (ETI). Doke (1983) first demonstrated that ROS members possibly function as the chemical signals required for hypersensitive response (HR) induction. The oxidative burst induced by pathogen-derived signals is essential to defense mechanisms throughout the plant (Fobert and Després, 2005). Apoplastic ROS are generated from plasma membrane-localized NADPH oxidases and cell wall-localized peroxidases. ROS are known as antimicrobial molecules involved in cell wall reinforcement and callose deposition. ROS also act as local and systemic secondary messengers triggering additional immune responses, including stomatal movement. Stomatal closure in response to various stress is brought about by loss of guard cell turgor, which is caused by ion channel modulation. A complex signaling network involving ROS production regulates these channels (Kim et al., 2010; Kollist et al., 2014). ASM treatment of the first leaves via fungal inoculation caused rapid H_2_O_2_ accumulation below the penetration site on ASM-untreated third leaves (Lin and Ishii, 2009; Park et al., 2019), which might have contributed to the triggered defense responses against fungal pathogen infection. Endogenous SA is essential for both apoplastic ROS production and stomatal closure induced by chitosan as microbe-associated molecular patterns (MAMPs) (Prodhan et al., 2017). Since ASM is a synthetic analog of SA, it is tempting to speculate that ASM induced stomatal closure is closely related to ROS production.

We demonstrate that ASM successfully suppresses disease development and bacterial multiplication in Japanese radish by activating stomatal defense. Furthermore, to gain knowledge into the ASM action mechanism on stomata, we investigated its effect on ROS production. ASM activated ROS production mediated by peroxidases, which in turn activated stomatal-based defense. Moreover, ASM also induces *PR* gene expression on Japanese radish. Importantly, ASM controlled bacterial blight disease not only on the treated leaves, but also on Japanese radish upper and lower leaves. Thus, our results highly support that ASM can be an additional disease management tool to prevent crop disease losses against bacterial pathogens.

## Materials and Methods

### Plant materials and chemical treatment

Japanese radish (*Raphanus sativus* var. *longipinnatus*) cv. Natsutsukasa plants were used for all experiments. Seedlings were grown in 9 cm pots at 24°C with a light intensity of 200 μmol/(m^2^ sec) and 16 h light/8 h dark in a growth chamber or were maintained in the greenhouse under a natural photoperiod at 21.2±2.4°C. Acibenzolar-S-methyl (ASM, marketed as ACTIGARD^®^) was supplied courtesy of Syngenta. To evaluate the effect of ASM on plant defense, Japanese radish fourth leaves were treated with ASM (100 ppm) by dip-treatment at 3 weeks after sowing or when fifth true leaves were unfolded. Salicylhydroxamic acid (SHAM) and diphenyleneiodonium chloride (DPI) were obtained from Sigma-Aldrich (Sigma-Aldrich, St Louis, MO, USA). Plants were treated with ASM 4 h, 1 d, and 1 w before *Pcal* inoculation (Supplementary Figure 1). Plants were dip-treated with water or mock-inoculated with water as controls.

### Bacterial strains and growth conditions

Pathogenic *Pseudomonas cannabina* pv. *alisalensis* strain KB211 (*Pcal*) (Sakata et al., 2019) was kindly provided by the Nagano Vegetable and Ornamental Crops Experiment Station, Nagano, Japan. *Pcal* was grown at 28°C for 24 h on Kings B (KB) (King et al., 1954) agar medium. For inoculum, bacteria were suspended in sterile distilled H_2_O, and cell densities measured at 600 nm (OD_600_) using a Biowave CO8000 Cell Density Meter (biochrom, Cambridge, UK).

### Bacterial inoculation

Plants were spray-inoculated with a bacterial suspension in sterile distilled water containing 0.025% Silwet L-77 (bio medical science, Tokyo, Japan). For *Pcal* lesion area and disease symptoms measurement, Japanese radish plants were spray-inoculated with a bacterial suspension (5 × 10^7^ colony forming units [CFU]/ml) to runoff. Dip-inoculation method was used for stomatal assay. For bacterial growth assay, Japanese radish plants were spray-inoculated with a bacterial suspension (5 × 10^7^ CFU/ml). The plants were then incubated in growth chambers at approximately 100% RH for the first 24 h, then at approximately 70% RH for the rest of the experiment. Lesion area and bacterial population were assessed 1-week post-inoculation (wpi).

### Real-time quantitative RT-PCR

For expression profiles of Japanese radish defense genes in response to ASM, plants were dip-treated with water as a control or ASM on fourth leaves. After 4 h total RNA was extracted using RNAiso Plus (Takara Bio, Shiga, Japan) according to the manufacturer’s protocol and used for real-time quantitative RT-PCR (qRT-PCR) as described (Ishiga and Ichinose 2016). Two micrograms of total RNA were treated with gDNA Remover (TOYOBO, Osaka, Japan) to eliminate genomic DNA, and the DNase-treated RNA was reverse transcribed using the ReverTra Ace^®^ qPCR RT Kit (TOYOBO). The cDNA (1:50) was then used for RT-qPCR with the primers shown below with THUNDERBIRD^®^ qPCR Mix (TOYOBO) on a Thermal Cycler Dice Real Time System (Takara Bio). Radish *Glyceraldehyde 3-phosphate dehydrogenase* (*GAPDH*) gene was used as an internal control. Average CT values, calculated using the second derivative maximum method from triplicate samples, were used to determine the fold expression relative to the controls (0 time). Primers used in gene-specific PCR amplification for *PR1* were 5′- AAAGCTACGCCGACCGACTACGAG -3′ and 5′- CCAGAAAAGTCGGCGCTACTCCA -3′; for *PR2*, 5′- GTACGCTCTGTTCAAACCGACCC -3′ and 5′- TTTCCAACGATCCTCCGCCTGA -3′; for *PR3*, 5′- TCTTTGGTCAGACTTCCCACGAG -3′ and 5′- GATGGCTCTTCCACACTGTCCGTA -3′; and for *GAPDH*, 5′- CGCTTCCTTCAACATCATTCCCA -3′ and 5′- TCAGATTCCTCCTTGATAGCCTT -3′.

### Stomatal assay

A modified method was used to assess stomatal response as described previously (Chitrakar and Melotto, 2010). Briefly, Japanese radish plants were grown for 3 weeks after sowing as described previously. *Pcal* was grown at 28°C for 24 h on KB agar medium, then suspended in sterile distilled water to 1 × 10^8^ CFU/ml. Dip-inoculated or water mock-inoculated radish leaves were directly imaged at 4 h post-inoculation (hpi) using a Nikon optical microscope (Eclipse 80i). Additionally, 1 week after ASM treatment, leaves were treated with SHAM (1 mM) and DPI (10 μM) with and without *Pcal*. The aperture width of at least 100 stomata was measured 4 h post inoculation and treatment (Supplementary Figure 1). The average and standard error for the stomatal aperture width were calculated. The stomatal apertures were evaluated in three independent experiments.

## Results

### ASM suppresses *Pcal* disease development in radish and triggers SAR

To investigate if ASM effectively suppresses *Pcal* disease development in Japanese radish plants, plants were inoculated with *Pcal*, 4 h, 1 d, and 1 w after ASM dip-treatment on only the fourth leaf. Then disease symptoms were observed 7 dpi (Supplementary Figure 1). ASM-treated Japanese radish plants showed less disease symptoms than control water-treated plants inoculated with *Pcal* in all treatments (Supplementary Figure 2). We also evaluated the disease development by measuring lesion areas. *Pcal* lesion area on ASM-treated Japanese radish plants was significantly smaller than that of control water-treated plants within 4 h post-treatment (hpt) (Figures 1A, B, C). Furthermore, lesion area was reduced not only on the treated Japanese radish leaves, but also on the untreated upper and lower leaves (Figures 1A, B, C). To investigate whether ASM suppresses bacterial growth as well as lesion area, we also measured bacterial population 7 dpi. In ASM-treated Japanese radish fourth leaves, *Pcal* populations were around ten times less compared to those of the water-treated control 4 hpt (Figure 2A). In addition, at 1 d post-treatment (dpt) and 1w post-treatment (wpt) with ASM, *Pcal* populations were around 100 times less compared to those of the water-treated control (Figures 2B, C). Moreover, in third and fifth leaves, *Pcal* populations were also less compared to those of the water-treated control (Figures 2A, B, C). These results indicate that ASM is able to suppress symptom development and bacterial multiplication in Japanese radish, and this effect is acquired systemically.

**Figure 1.**
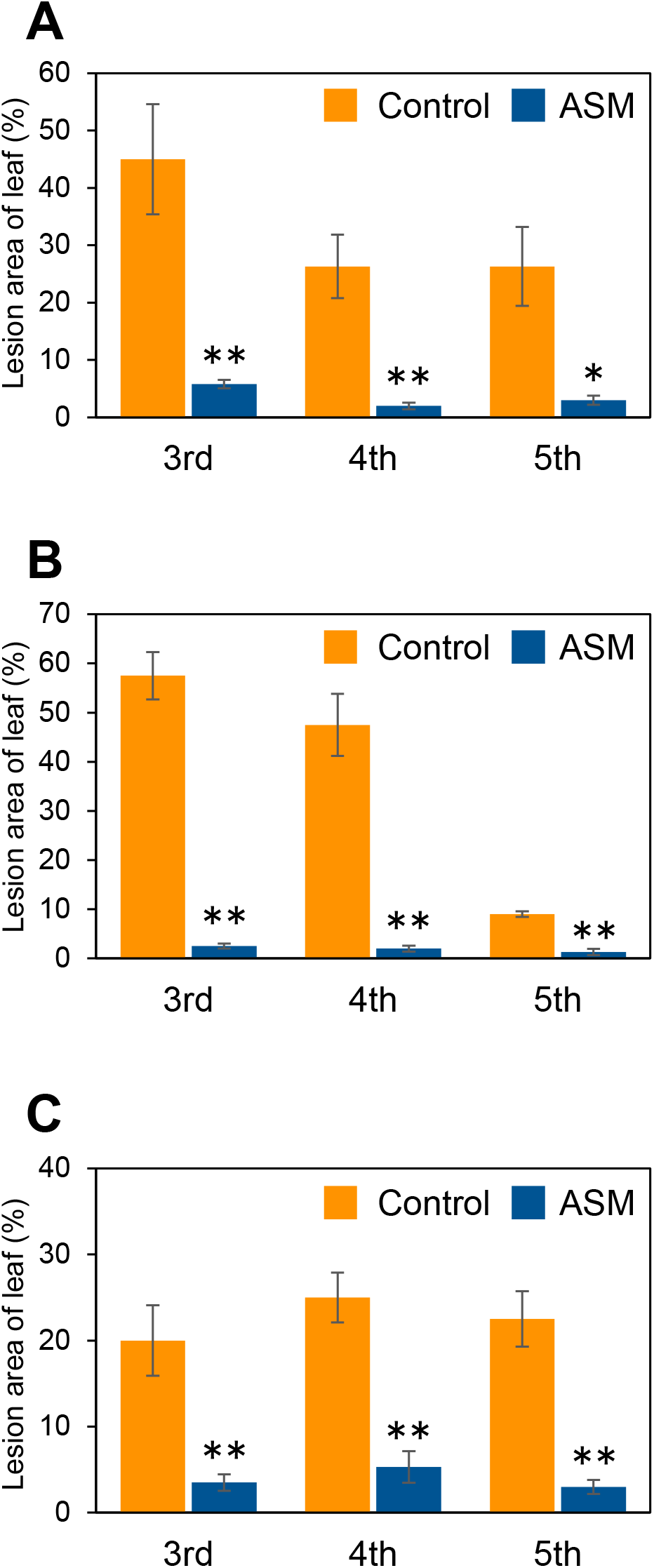
Lesion area (%) of Japanese radish inoculated with *Pcal* after dip-treatment with ASM on fourth leaves. Greenhouse grown Japanese radish plants were spray-inoculated with *Pcal* (5 × 10^7^ CFU/ml) 4 h **(A)**, 1 d **(B)**, and 1 w **(C)** after ASM dip-treatment (100 ppm) on fourth leaves. Disease symptoms were monitored by measuring lesion areas at 1-week post inoculation (wpi). Vertical bars indicate the standard error for four independent replications. Asterisks indicate a significant difference from the water treatment control in a *t*-test (\**p* < 0.05; ** *p* < 0.01).

**Figure 2.**
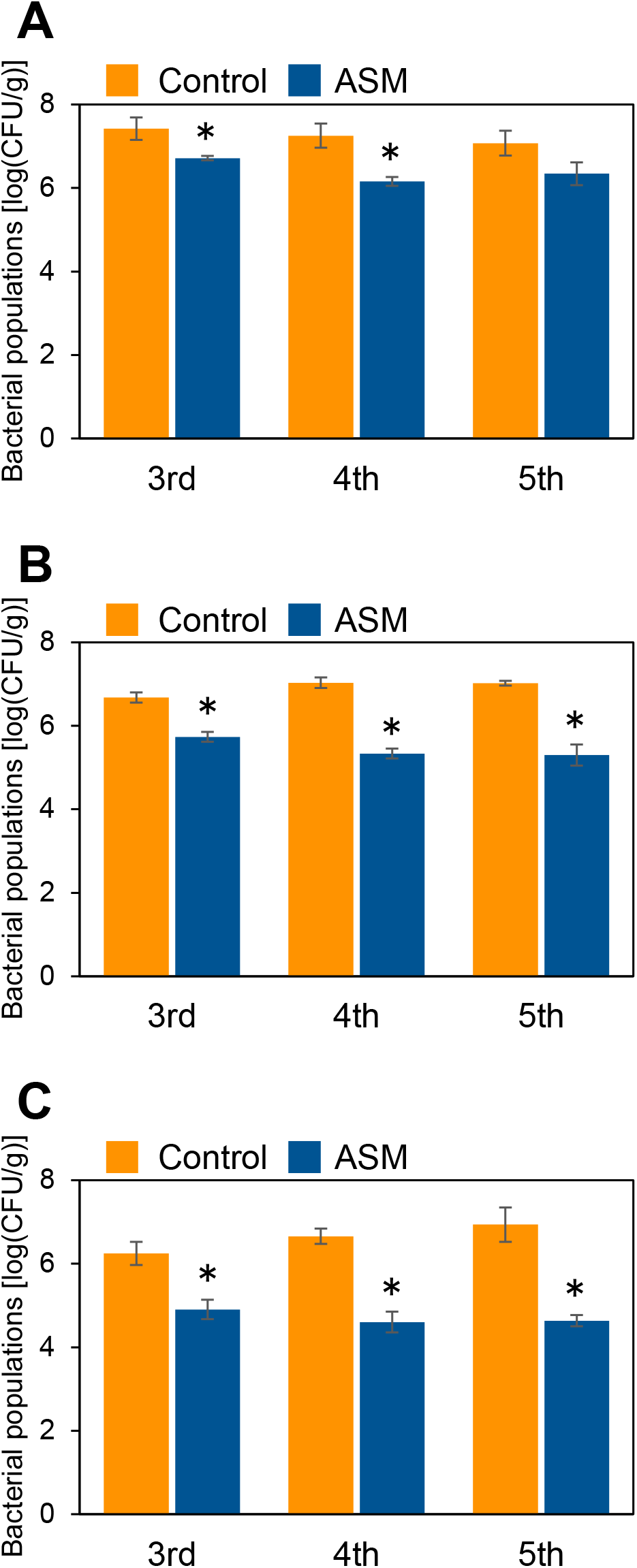
Bacterial populations of *Pcal* inoculated plants after ASM dip-treatment. Three-week-old Japanese radish plants were spray-inoculated with *Pcal* suspensions (5 × 10^7^ CFU/ml) 4 h **(A)**, 1 d **(B)**, and 1 w **(C)** after ASM dip-treatment (100 ppm) on fourth leaves. Bacterial populations were determined by dilution plating on selective medium as described in the methods section at 1-week post inoculation (wpi). Vertical bars indicate the standard error for three independent experiments. Asterisks indicate a significant difference from the water treatment control in a *t*-test (\**p* < 0.05; ** *p* < 0.01).

### ASM activates stomatal-based defense against *Pcal*

We demonstrated that ASM activated stomatal-based defense against *Pcal*, resulting in reduced disease development and bacterial populations on cabbage (Ishiga et al., 2020; Ishiga et al., unpublished). To examine whether ASM activates stomatal-based defense on Japanese radish, we observed stomatal response after *Pcal* inoculation after ASM dip-treatment on fourth leaves. After *Pcal* inoculation, ASM treated Japanese radish leaves exhibited stomatal closure (Figures 3B, E, H). Interestingly, we also observed stomatal closure of the untreated upper and lower leaves (Figures 3A, C, D, F, G, I). These results indicate ASM activated stomatal-based defense against *Pcal* on radish systemically. Furthermore, stomatal closure was observed on ASM mock-inoculated Japanese radish leaves at 1 dpt and 1 wpt (Figures 3D, E, F, G, H, I), but not 4 hpt (Figures 3A, B, C). These results indicate that ASM suppresses stomatal opening regardless of *Pcal* inoculation 1 dpt.

**Figure 3.**
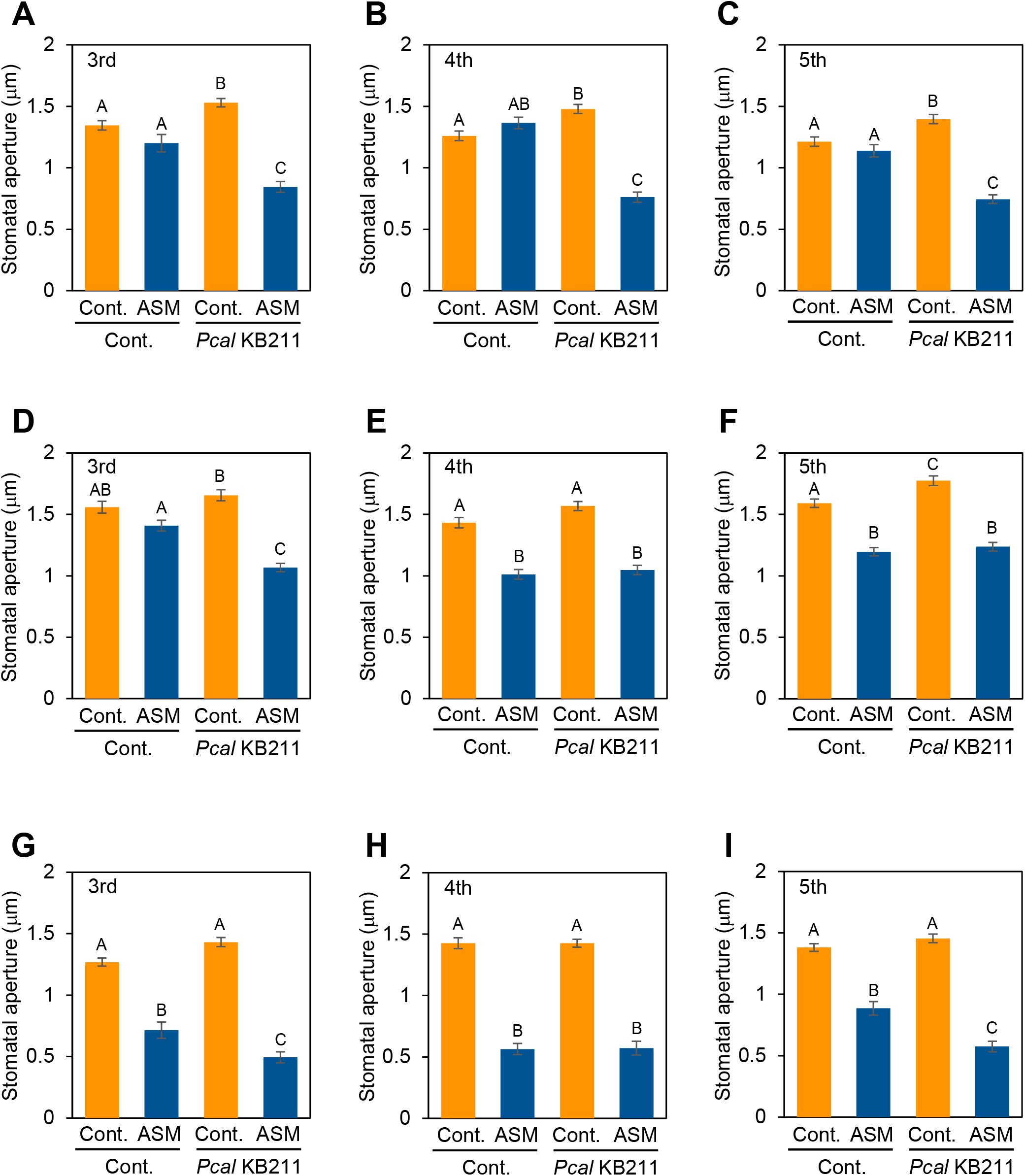
Stomatal aperture width (μm) in Japanese radish plants dip inoculated with *Pcal* suspensions (1× 10^8^ CFU/ml) after ASM treatment. Japanese radish leaves were inoculated with *Pcal* 4 h **(A–C)**, 1 d **(D–F)**, and 1 w **(G–I)** after ASM dip-treatment (100 ppm) on fourth leaves. Stomatal aperture width (μm) was measured in third (**A, D, G**), fourth **(B, E, H)**, and fifth **(C, F, I)** leaves. Each time point after ASM treatment, radish leaves were dip-inoculated, then imaged using a Nikon optical microscope (Eclipse 80i). In all bar graphs, vertical bars indicate the standard error for three biological replicates. Significant differences (*p* < 0.05) are indicated by different letters based on a Tukey’s honestly significant difference (HSD) test.

### ASM induced stomatal closure by inducing peroxidase-dependent ROS production

Reactive oxygen species (ROS) are essential second messengers in stomatal immunity (Joshi-Saha et al., 2011). Stomatal closure by the phytohormone salicylic acid (SA) requires ROS production by cell wall peroxidases (Khokon et al., 2011; Mori et al., 2001). Since ASM is a synthetic analog of SA, we investigated whether ASM induced stomatal closure by producing ROS through peroxidase by using a peroxidase inhibitor, salicylhydroxamic acid (SHAM), and an NADPH oxidase inhibitor, diphenylene iodonium (DPI). ASM-induced stomatal closure was suppressed by SHAM (Figures 4A, B, C), indicating that cell wall peroxidases are involved in mediating ROS production during ASM-induced stomatal closure. Conversely, exogenous DPI application did not inhibit the ASM-induced stomatal closure (Figures 4A, B, C). We obtained the same results after *Pcal* inoculation (Supplementary Figures 3A, B, C). These results indicate that ASM induces stomatal closure via ROS production mediated by peroxidases.

**Figure 4.**
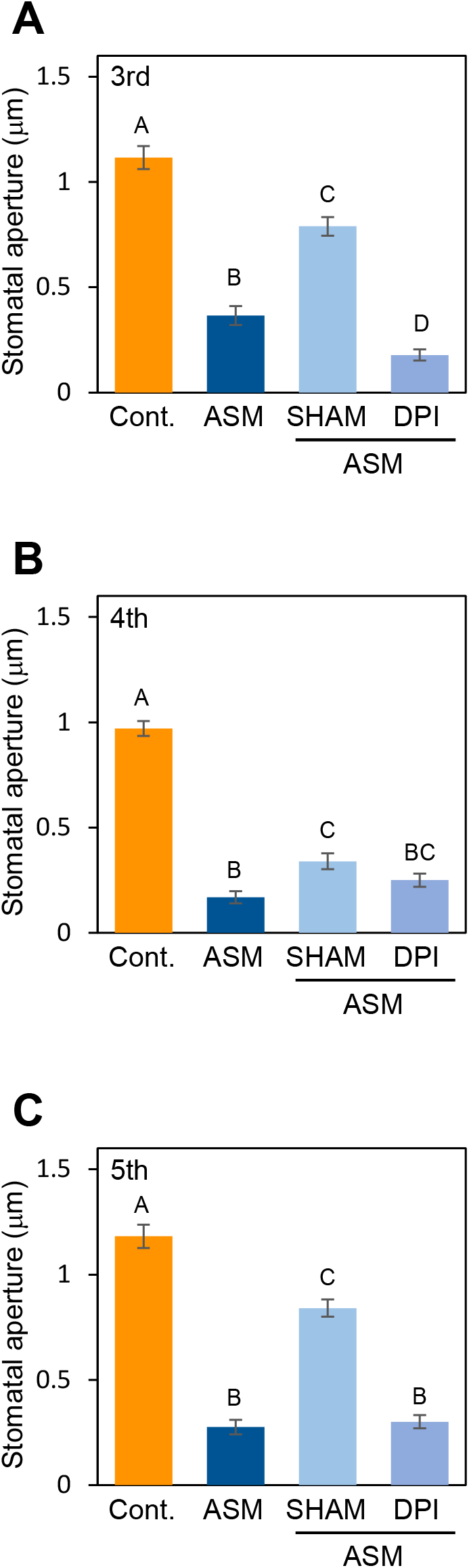
Stomatal aperture width (μm) in Japanese radish plants treated with SHAM and DPI after ASM treatment. Japanese radish leaves were treated with SHAM (1 mM) and DPI (10 μM) 1 w after ASM dip-treatment (100 ppm) on fourth leaves. Stomatal aperture width (μm) was measured on third **(A)**, fourth **(B)**, and fifth **(C)** leaves using a Nikon optical microscope (Eclipse 80i). In all bar graphs, vertical bars indicate the standard error for three biological replicates. Significant differences (*p* < 0.05) are indicated by different letters based on a Tukey’s honestly significant difference (HSD) test.

### ASM induces defense-related gene expression in Japanese radish

ASM also induced defense-related gene expression, including PR proteins (Ishiga et al., 2020; Kouzai, et al., 2018; Kouzai, et al., 2018; Nakashita et al., 2002; Yasuda et al., 2008). To investigate the effect of ASM treatment on Japanese radish defense gene expression, we investigated the expression profiles of *PR1*, *PR2*, and *PR3* in response to ASM. The expression of *PR1*, *PR2*, and *PR3* was significantly induced in fourth leaves 4 h after ASM treatment (Figures 5A, B, C). Furthermore, in both untreated upper and lower leaves, all gene expression was induced or tended to be induced (Figures 5A, B, C). These results indicate that ASM induced defense-related gene expression triggered systemic acquired resistance in Japanese radish.

**Figure 5.**
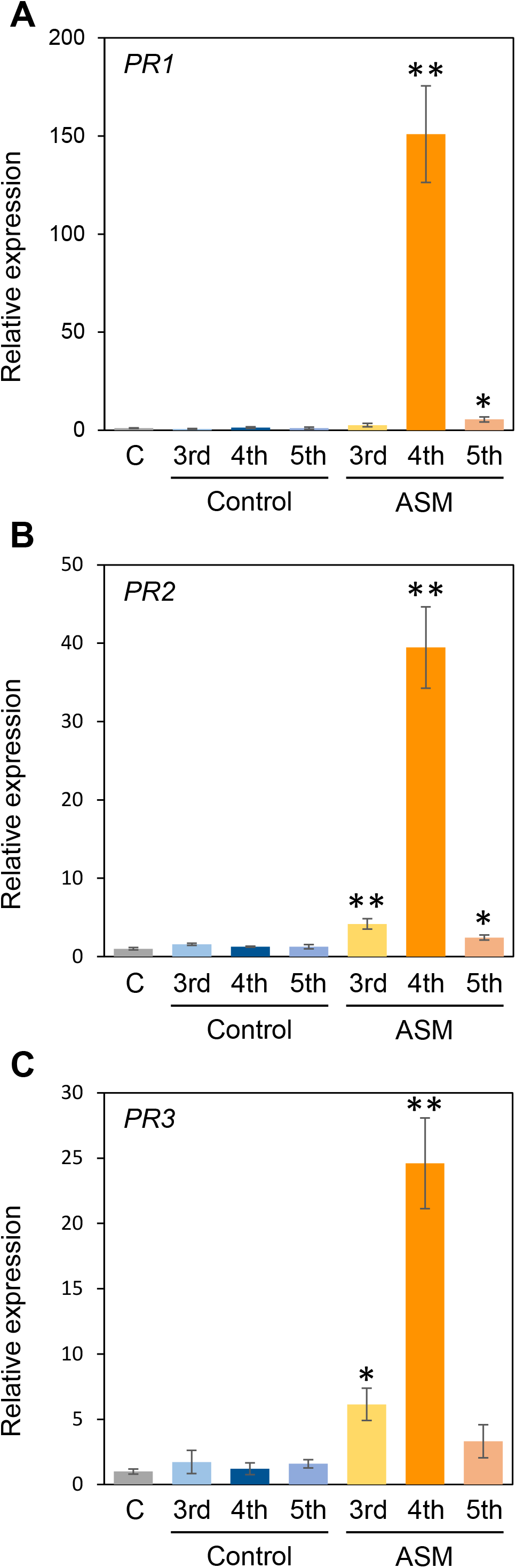
Gene expression profiles involved in Japanese radish plant defense after ASM treatment. Three-week-old Japanese radish plants were treated with water as a control and ASM (100 ppm). After 4 h expression of *PR1* **(A)**, *PR2* **(B)**, and *PR3* **(C)** was determined using real-time quantitative reverse transcription-polymerase chain reaction (RT-qPCR) with gene-specific primer sets. Expression was normalized using *GAPDH*. Vertical bars indicate the standard error for three biological replicates. Asterisks indicate a significant difference from the water control in a *t*-test (\**p* < 0.05; ** *p* < 0.01).

## Discussion

There is an increased demand for developing sustainable disease control strategies. ASM is predicted as an additional disease management tool to protect numerous crops from pathogens without the occurrence of chemically resistant strains. Our results clearly showed that an ASM dip-treatment effectively suppressed *Pcal* lesion formation and bacterial multiplication on Japanese radish plants (Figure 1; Figure 2; Supplementary Figure 2). Previously, we also demonstrated that an ASM soil drench suppressed *Pcal* disease development associated with reduced bacterial population in cabbage (Ishiga et al., 2020). These results indicate that ASM is a powerful tool against *Pcal*, which is the causal agent of bacterial blight of cruciferous plants.

The most important finding was that ASM greatly enhanced the systemic activation of defense response. The reduction of disease development and bacterial growth was observed not only on treated fourth leaves, but also on untreated third and fifth leaves (Figure 1; Figure 2; Supplementary Figure 2). Additionally, ASM triggered stomatal-based defense and SAR-related gene expression induction on both treated and untreated leaves (Figure 3; Figure 5). Cools and Ishii (2002) demonstrated that ASM pre-treatment of cucumber first leaves protected whole plants from infection with the virulent fungal pathogen *Colletotrichum orbiculare*. Three hours post-ASM treatment on first leaves, ASM-induced systemic priming of *PAL1* expression in cucumber plants occurred rapidly, with enhanced expression in the third leaves (Cools and Ishii, 2002). Interestingly, our results showed ASM-induced systemic defense was acquired not only on untreated upper leaves, but also on untreated lower leaves (Figure 5). Kubota and Abiko (2000) demonstrated that 7 days after a first inoculation with *Colletotrichum lagenarium* and *Pythium ultimum* on cucumber cotyledons and primary leaves, induced resistance was observed against the challenge inoculation with those pathogenic fungi in hypocotyl. These results implied that foliar leaf treatment can prevent disease of lower plant parts, including hypocotyls and roots (Kubota and Abiko, 2000). Importantly, *Pcal* causes discoloration of Japanese radish hypocotyl and root (Takikawa and Takahashi, 2014). It is expected that ASM-induced systemic defense could be a powerful disease management tool against bacterial blight on Japanese radish.

The gene expression profiles of *PR1*, *PR2*, and *PR3* were induced within 4 h in both treated and untreated leaves (Figure 5). Benzo (1,2,3) thiadiazole-7-carboxylic acid (acibenzolar), but not ASM itself was detected in untreated third leaves 3 dpt on first leaves (Ishii et al., 2002). SABP2 catalyzes ASM conversion into acibenzolar to induce SAR (Tripathi et al., 2010). Therefore, acibenzolar may function similar to SA to activate NPR1 and SAR. Although the ASM control effect was obvious within 4 hpt (Figure 1; Figure 2) and SAR-related gene expression was also induced within 4 hpt (Figure 5), acibenzolar was not detected in untreated leaves 1 dpt (Ishii et al., 2002). Therefore, it is tempting to speculate that putative signal molecules are rapidly translocated from treated to untreated leaves. Several candidates for this long-distance signal have been identified, including methyl salicylate (MeSA), an SFD1/GLY1-derived G3P-dependent signal, the lipid-transfer protein DIR1, the dicarboxylic acid azelaic acid (AzA), the abietane diterpenoid dehydroabirtinal (DA), jasmonic acid (JA), and the amino acid derivative pipecolic acid (Pip) (Dempsey and Klessig, 2012). Among them, Pip acts as primary regulator of SAR (Návarová et al., 2012; Zeier, 2013). In *Arabidopsis thaliana*, these responses include camalexin (phytotoxin) accumulation and expression of defense-related genes, including *ALD1*, *FMO1*, and *PR1* (Bernsdorff et al., 2016). Further analysis will be needed to elucidate the mobile signals which induce SAR in Japanese radish.

ASM induced stomatal-based defense against *Pcal* within 4 hpt in both cabbage and Japanese radish (Figure 3; Ishiga et al., 2020). We also showed that ASM-triggered stomatal closure was suppressed by a peroxidase inhibitor, SHAM but not by an NADPH oxidase inhibitor, DPI (Figure 4; Supplementary Figure 3). Khokon et al. (2011) reported that SA-induced stomatal closure accompanied ROS production mediated by peroxidase in *A. thaliana*. Therefore, these results indicated that ASM-induced stomatal closure is closely related to ROS production through peroxidase. Our results also showed that ASM induced stomatal-based defense was observed not only on treated leaves, but also on untreated leaves (Figure 3). Lin and Ishii (2009) demonstrated that rapid H_2_O_2_ accumulation occurred in cucumber stem xylem fluids during ASM-induced SAR.

Takahashi and Shinozaki (2019) reported that light stress-mediated ABA accumulation induced the ROS/Ca^2+^ wave propagates from local leaves to systemic leaves with a rapid transmission speed, suggesting that local ROS/Ca^2+^ wave functions as long-distance signal to activate stomatal closure during light stress conditions. Therefore, it is possible that ROS induced by ASM treatment activates stomatal closure in a systemic response during pathogen infection.

Based on our results, we propose a stomatal closure model induced by ASM treatment (Figure 6). ASM induces ROS production mediated by SHAM-sensitive peroxidase, leading to stomatal closure. Then, local ROS functions as a long-distance signal to induce stomatal closure in a systemic response after ASM treatment. In addition to ROS, it is expected that some mobile signals are rapidly transported and induce ROS production, and in turn stomatal closure. Further analysis will be needed to understand the ASM-induced SAR mechanism.

**Figure 6.**
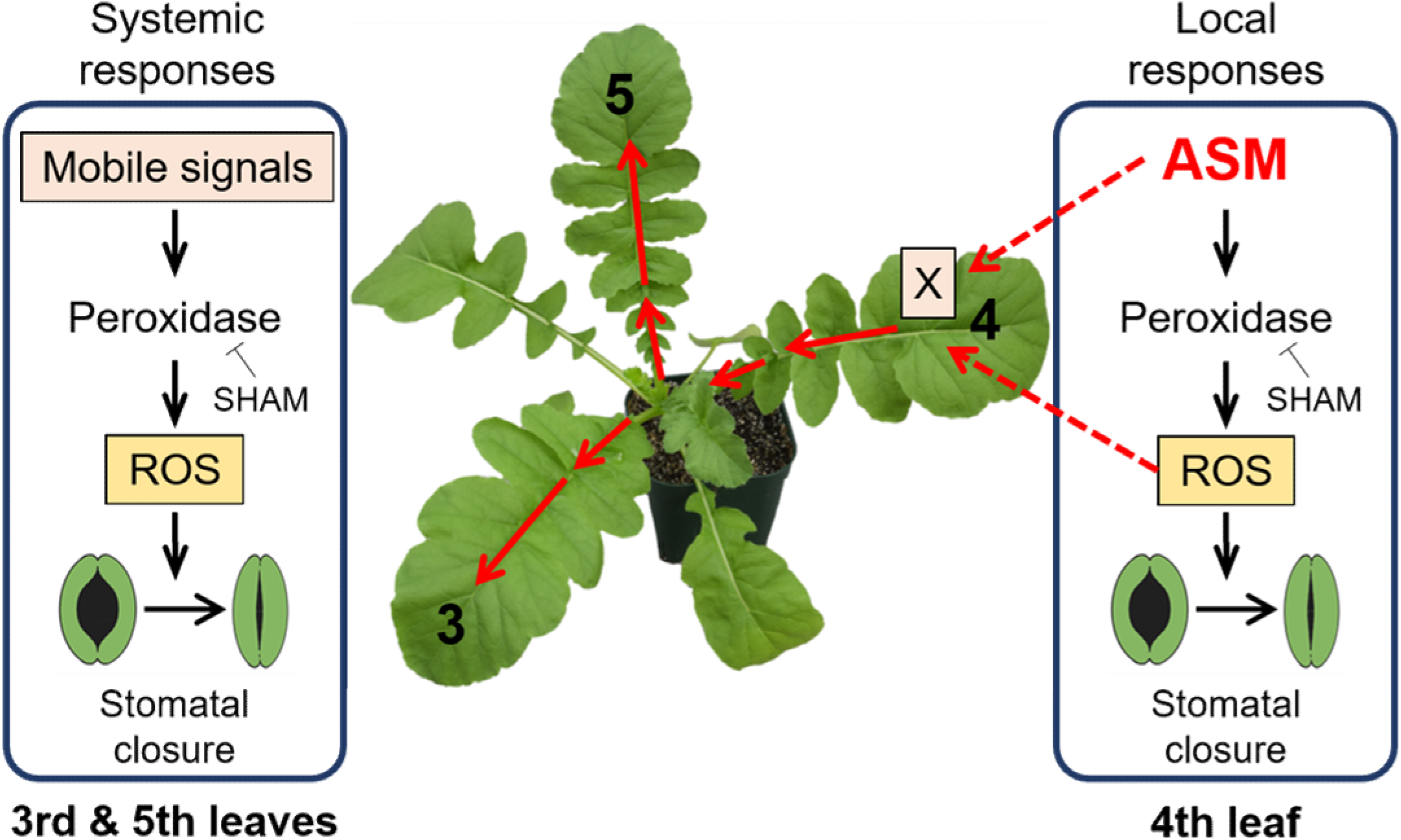
Proposed model of stomatal closure induced by ASM treatment by inducing peroxidase-dependent ROS production. ASM induces stomatal closure by inducing ROS production through peroxidase. Then, local ROS and some mobile signals function as long-distance signals to induce stomatal closure in a systemic response after ASM treatment. X indicates mobile signals.

We clearly demonstrated that ASM dip-treatment protects Japanese radish plants against *Pcal*, a causal agent of bacterial blight disease, by activating SAR, such as stomatal-based defense. Furthermore, we observed that ASM induced defense response was acquired not only on treated leaves, but also untreated upper and lower leaves. These results highlight the role of ASM as an efficient sustainable disease control strategy.

## Acknowledgments

We thank Dr. Christina Baker for editing the manuscript. *Pcal* was kindly provided by the Nagano Vegetable and Ornamental Crops Experiment Station, Nagano, Japan. This work was supported in part by the JST ERATO NOMURA Microbial Community Control Project, JST, Japan.

