## Supplemental Figures for "Acibenzolar-S-methyl activates stomatal-based defense systemically in Japanese radish by inducing peroxidase-dependent reactive oxygen species production"

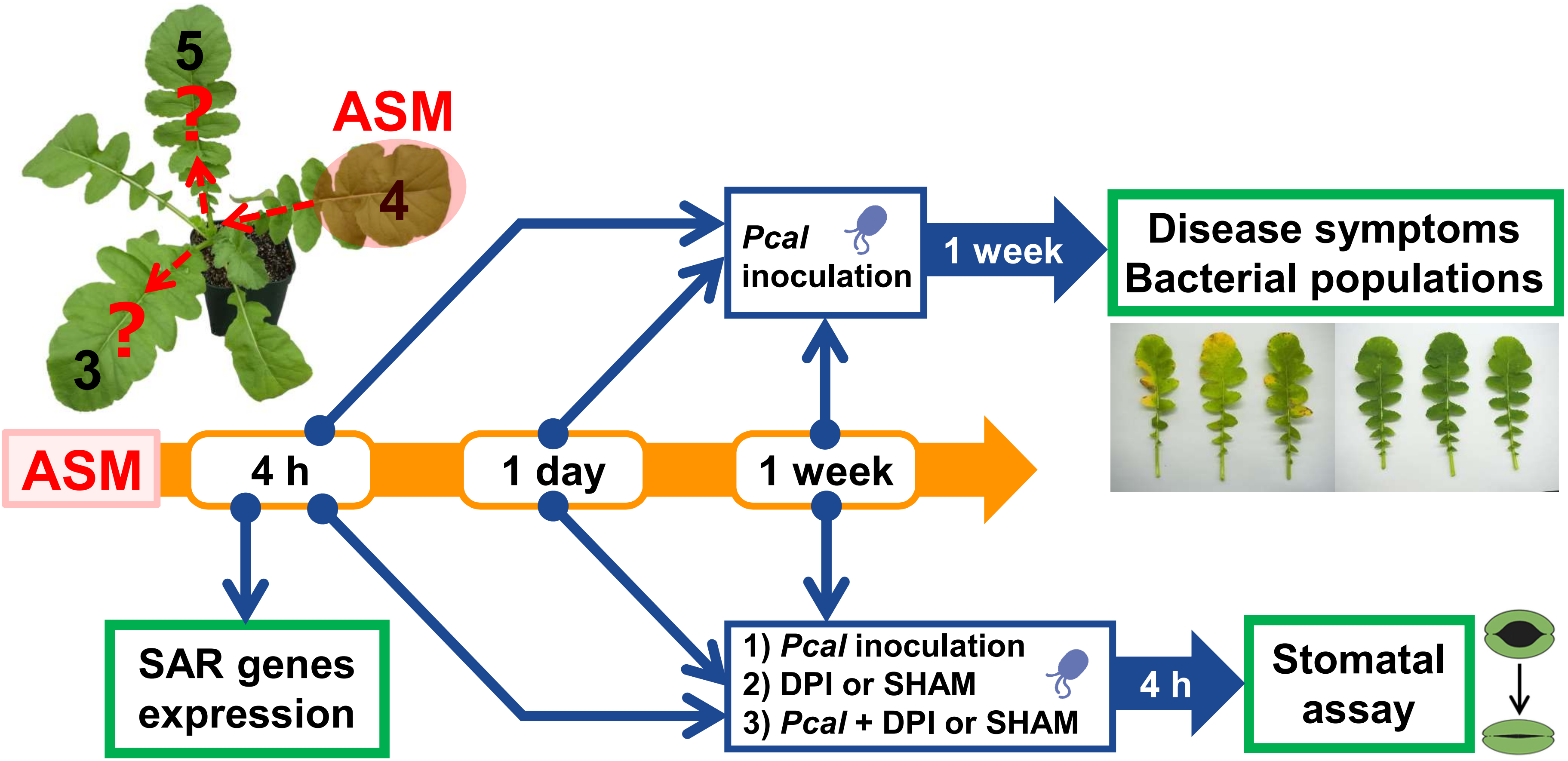


**Supplementary Figure 1. The workflow of this study.** Japanese radish plants were inoculated with *Pcal* 4 h, 1 d, and 1 w after ASM dip-treatment on only the fourth leaf. Disease symptoms (area and bacterial population) were assessed at 1-week post-inoculation (wpi). For stomatal assay, plants were 1) inoculated with *Pcal*, 2) treated with DPI and SHAM, and 3) inoculated with *Pcal* and treated with DPI and SHAM, 4 h, 1 d, and 1 w after ASM dip-treatment on only fourth leaves. For SAR genes expression profiles, plants were dip-treated with ASM on fourth leaves. Total RNAs were extracted 4 h after dip-treatment with ASM on fourth leaves and then used for RT-qPCR.

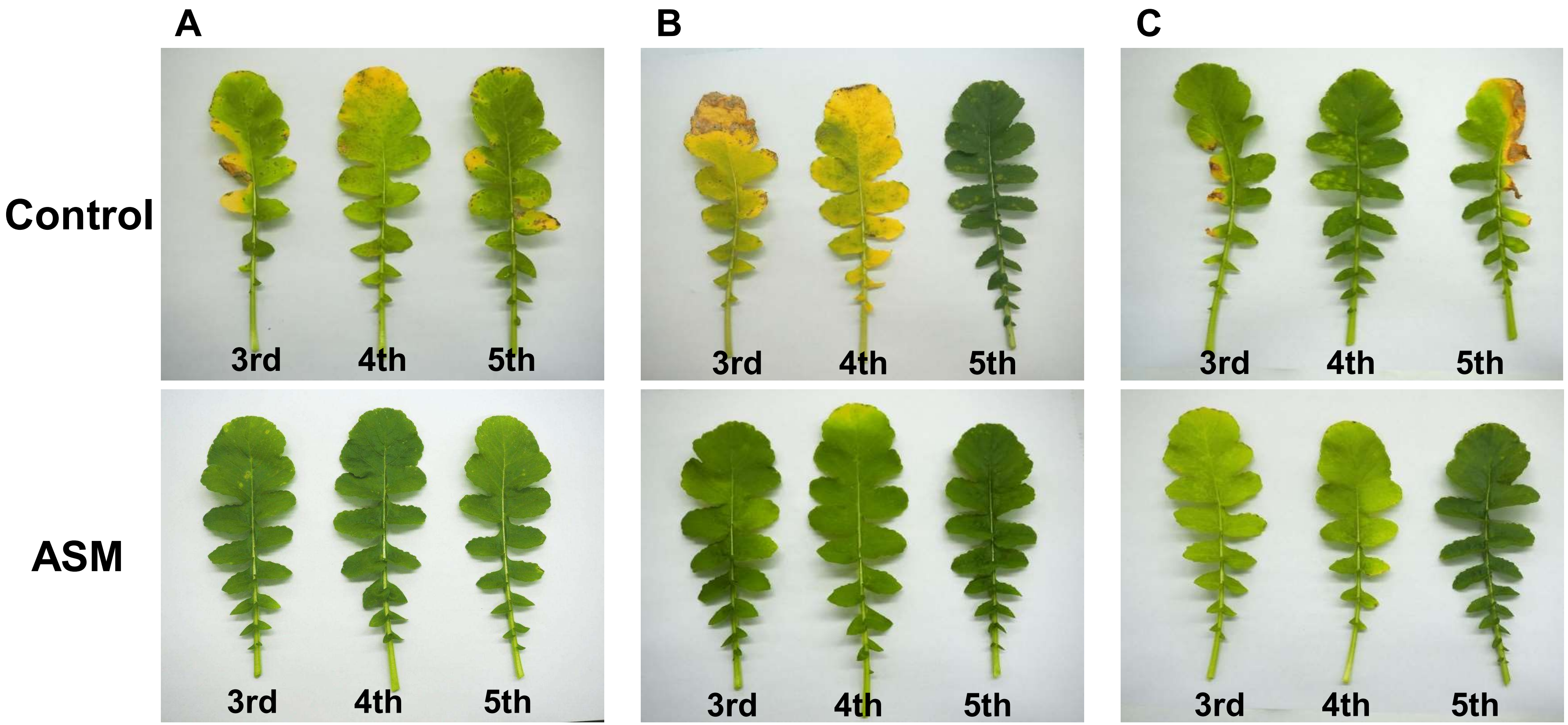


**Supplementary Figure 2. Symptom development of Japanese radish inoculated with *Pcal* after the dip-treatment with ASM on fourth leaves.** Greenhouse grown Japanese radish plants were spray-inoculated with *Pcal* ($5 \times 10^7$ CFU/ml) 4 h **(A)**, 1 d **(B)**, and 1 w **(C)** after ASM dip-treatment (100 ppm) on fourth leaves. Disease symptoms were observed 7 days post-inoculation.

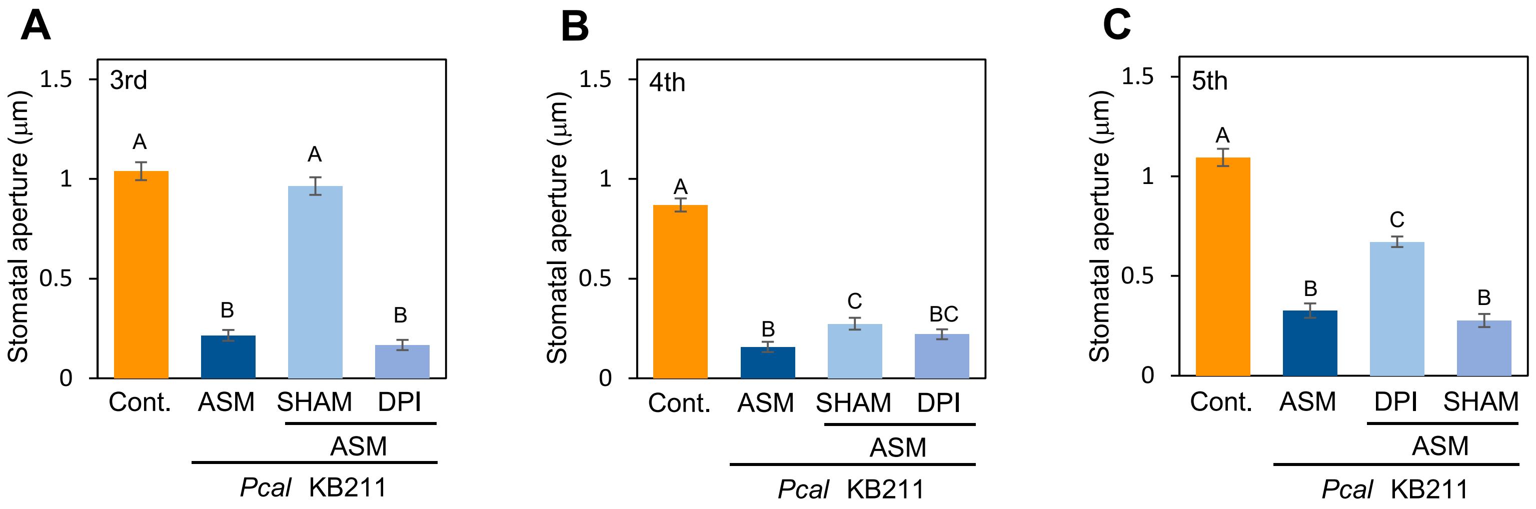


**Supplementary Figure 3. Stomatal aperture width (µm) in Japanese radish plants dip-inoculated with *Pcal* suspensions (1 × 10$^{8}$ CFU/ml) and treated with SHAM and DPI after ASM treatment.** Japanese radish leaves were inoculated with *Pcal* and treated with SHAM (1mM) and DPI (10 µM) after ASM dip-treatment (100 ppm) on forth leaves. Stomatal aperture width (µm) was measured on third **(A)**, fourth **(B)**, and fifth **(C)** leaves using a Nikon optical microscope. In all bar graphs, vertical bars indicate the standard error for three biological replicates. Significant differences ($p < 0.05$) are indicated by different letters based on a Tukey's honestly significant difference (HSD) test.
